# An atlas of TF driven gene programs across human cells

**DOI:** 10.1101/2025.05.30.657075

**Authors:** J. Patrick Pett, Martin Prete, Duy Pham, Nick England, Hao Yuan, Elena Prigmore, Liz Tuck, Agnes Oszlanczi, Ken To, Chuan Xu, Chenqu Suo, Emma Dann, Peng He, Veronika Kedlian, Kazumasa Kanemaru, James Cranley, Ling Yang, Rasa Elmentaite, Amanda J. Oliver, Ana-Maria Cujba, Batuhan Cakir, Simon Murray, Krishnaa T. Mahbubani, Kourosh Saeb-Parsy, Laure Gambardella, Maria Kasper, Muzlifah Haniffa, Martijn C. Nawijn, Sarah A. Teichmann, Kerstin B. Meyer

## Abstract

Combinations of transcription factors (TFs) regulate gene expression and determine cell fate. Much effort has been devoted to understanding TF activity in different tissues and how tissue-specificity is achieved. However, ultimately gene regulation occurs at the single cell level and the recent explosion in the availability of single cell gene expression data now makes it possible to understand TF activity at this granular level of resolution.

Here, we leverage a large collection of Human Cell Atlas (HCA) single cell data to explore TF activity by examining cell-type and tissue-specific sets of target genes, or regulons. We compile a regulon atlas, CellRegulon, and map the activity of TFs in an extensive set of healthy adult and foetal tissues spanning hundreds of cell types. Using CellRegulon, we describe dynamic patterns of co-regulation, associate TF-modules with different cellular functions and characterise the distribution of active TFs and TF families across cell types. We show that CellRegulon can link disease gene expression signatures to cell types and TFs relevant to the disease. Finally, using a newly generated multiome dataset of the adult lung, we show how CellRegulon can be extended into an enhancer-gene regulatory network (eGRN) to improve cell-type associations with genetic risk loci for diseases, such as childhood onset asthma, COPD and IPF, and to identify high risk gene modules. Our database for easy download and interactive exploration allows researchers to understand key gene modules activated at cell type transitions and will therefore be valuable for tasks such as cell type engineering (https://www.cellregulondb.org).

## Introduction

The recent rise of single cell technologies and resources such as the Human Cell Atlas (HCA) allow recording gene expression and chromatin accessibility from a large variety of different cell types and conditions^1^. Leveraging these vast resources now opens up the possibility of characterizing cell-type-specific gene regulatory networks on a large scale at single cell resolution. This can be achieved by combining computational approaches with known molecular interaction preferences^2,3^.

Transcription factors (TFs) play a central role in determining the state of cells by governing transcriptional changes through gene regulatory networks. They bind to specific binding sites, often in promoters or enhancers to regulate the expression of their target genes. More than 1600 human TFs have been identified today and grouped into different families based on their DNA binding motifs^3,4^.

The activity of transcription factors is crucial for determining cellular fate and differentiation into distinct cell types. The Yamanaka factors can reprogram differentiated cells into pluripotent stem cells^5^ and, for example, a combination of three TFs was shown to convert fibroblasts into functional neurons *in vitro*^6^. Mutations in TFs *in vivo* have been linked to various monogenic diseases, such as loss of function mutations in HOX genes resulting in developmental conditions^7^. The role of TFs in the regulation of cellular functions has been studied in a large variety of tissues and the coordinated activity of multiple transcription factors is required for the differentiation of many cell types^8–10^.

Linking TFs to their target genes requires identifying the locations of TF binding sites, as well as matching those locations to target genes. Traditionally, transcription factor targets have been determined using ChIP-seq^11^ experiments, which identify binding sites across the genome for individual TFs. However, this approach is not easily scalable to larger numbers of TFs and in particular cell-type specific ChIP-seq data is available only for a limited number of primary cell types^12^. On the other hand, DNA binding specificities of TFs can be interrogated *in vitro* in high throughput e.g. using SELEX^13^ or PBM^14^ protocols.

Computational methods for the prediction of TF targets often leverage co-expression of TFs with their regulated targets^15^. However, co-expression does not always reflect a causal relationship. Therefore, methods like Scenic^2,16^ combine co-expression with the presence of DNA binding motifs to predict target genes for TFs, thereby increasing accuracy. As TFs often act in a cell type specific manner through epigenetic effects and interactions with other factors, cell type specific data is crucial for computational methods to delineate TF target-gene regulation and is becoming increasingly available in the form of single cell atlases.

Here, we identify sets of TF target genes, or regulons, resolved by cell type and tissue, compiled from a large number of single cell datasets, in total including 5 million cells covering 15 organ systems. This comprehensive single cell atlas of gene regulatory programs, CellRegulon, encompasses more than 700,000 regulons, offering a detailed census of TF dependent transcriptional states. We demonstrate that our atlas allows exploring TF activity across cell types and states, captures known cell state transitions and can be used to predict cell-type specific TF activity from bulk RNAseq data, for example to give insights into disease.

Furthermore, by integrating newly generated multiome data from the adult lung, we demonstrate that CellRegulon can be extended to enhancer-gene regulatory networks (eGRN) to improve cell-type associations with genetic risk loci for complex lung diseases including asthma, COPD and IPF. Our atlas is available for easy interactive queries through a web-interface and a dedicated python package for a wide range of applications.

## Results

### Census of transcription factors across single cell datasets

Bulk expression data has helped to delineate the roles of TFs across cell types in the past^17^. Large collections of single cell expression and chromatin accessibility data now allow a more fine-grained mapping of TF functions. Together with improved measurements of TF binding specificities^18^ and-sites, with methods such as CUT&RUN^19^, this will help to achieve an increasingly detailed characterization of human gene regulatory mechanisms.

To investigate TF function with much larger granularity we leveraged recently established single cell datasets to compute cell-type specific coregulated expression patterns of genes with common TF binding sites across a broad range of human tissues (Fig. 1a). To this end we gathered scRNA-seq datasets spanning 15 organ systems, including eye, oral cavity, endocrine-, reproductive-, respiratory-and musculoskeletal systems, kidney, bladder, vasculature, immune system, gastrointestinal tract as well as hepatobiliary system, heart and skin^20–30^. We used an adaption of the established pyScenic pipeline^2^, to compute cell type specific regulons taking account of co-expression of TF and target genes as well as the presence of known TF binding motifs in target gene promoters. In total, we inferred regulons for around 550 cell types and 1300 transcription factors (TFs) across multiple tissues and assembled a database of cell type specific, annotated regulons representing gene regulatory interactions across diverse human cell types.

**Figure 1:**
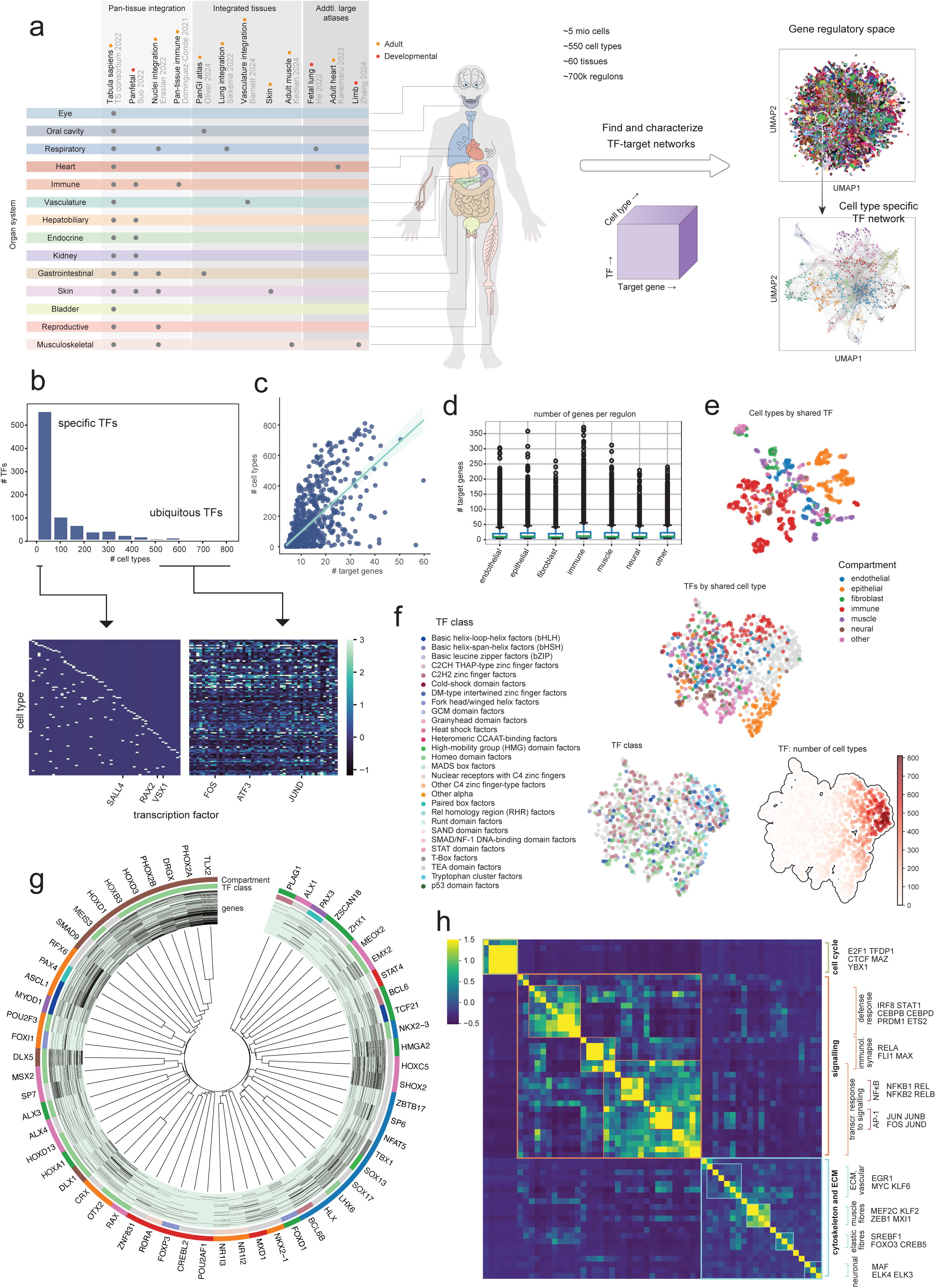
Regulon atlas overview. **(A)** Overview of included studies, spanning 14 organ systems. Studies include large single cell atlases integrated across and per-tissue. In total over 5 million cells have been processed to infer regulons for 550 cell types and 60 tissues. Right: TFs form a densely connected network that can be further explored on a cell type basis. **(B)** Histogram showing number of TFs per number of cell types. Most TFs are specific for a small number of cell types, while ubiquitous TFs account for most of the regulons with varying target genes across cell types and tissues. A TF was defined as specific or ubiquitous if it was found in <5 or>500 cell types, respectively. **(C)** Relationship between the number of target genes and number of cell types per TF. Ubiquitous TFs tend to target a larger number of genes across cell types. **(D)** Number of genes per regulon shown per cell compartment. Across compartments, most regulons target below 20 genes, while some target several hundreds. **(E)** Similarity of cell types by shared TFs. UMAP with cell types as points and Jaccard coefficients of common TFs as a similarity metric. Cell types from the same compartment often share TFs. **(F)** Similarity of transcription factors by shared cell types. UMAP with TFs as points and Jaccard coefficients of common cell types as a similarity metric. Clusters are less distinct than in (E). Top and right: For non-ubiquitous TFs, cell type and compartment have been assigned based on majority-voting. Some groups of similar TFs associate with the same cell compartment, while others are mixed. Left: Some TF groups also show enrichment by TF class, although more broadly than for cell types. **(G)** Specific TF clustering. Top-specific TFs are shown, clustered by shared target genes. They form groups belonging to the same cell type compartment and less often the same TF class. **(H)** Ubiquitous TF clustering. Top ubiquitous TFs are clustered by shared target genes and annotated for the functions of regulated genes. They regulate broad functions, such as cell cycle, cell signalling and cytoskeleton related.

When visualised in a low-dimensional embedding, the space of all ∼700k regulons presents in a spherical shape, reflecting the highly interconnected nature of TF regulation^31^ with varying combinations of TFs across cell types (Fig. 1a, right top). For subsets of the whole dataset, however, clusters of TFs regulating similar sets of target genes can be observed (Fig. 1a, right bottom). Examining the presence of regulons for all TFs across all cell types reveals a scale-free distribution (Fig. 1b), in which many TFs drive gene expression in only a small number of cell types specifically, while a smaller number of TFs are found across most cell types. Such ubiquitous TFs included TFs involved in general functions like proliferation, stress response and metabolism (*FOS*, *JUND*, *ATF3*), while specific TFs were associated with more specialised roles such as eye development (*VSX*, *RAX2*).

Ubiquitous TFs that regulate genes in most cell types tend to have larger numbers of target genes (Fig, 1c) with a varying composition across cell types. However, TFs with a large number of targets are found in both cell-type specific and ubiquitous ranges (Fig. 1c). Most TFs are predicted to regulate only around 20 target genes, while some are predicted to regulate hundreds of genes (Fig. 1d). Similarly, most genes are regulated by only a few TFs, while some are predicted to be regulated by several dozen (Extended Fig. 1a).

Plotting cell types as points in an embedding based on their similarity in terms of shared TF usage (Jaccard coefficient) shows cell types of the same compartment clustering together, with large groups formed by immune and epithelial cells (Fig. 1e), likely reflecting similar gene programmes that are active in cells from the same compartment (see below). Plotting TFs as points in an embedding based on their shared presence in cell types (Fig. 1f) shows a less distinct clustering compared to the cell type embedding (Fig. 1e), highlighting that TFs can be active across a large range of cell types, where they are likely to have distinct

functions. Nevertheless, distinct classes of TFs (based on structural homology) that are not ubiquitous (Fig. f) show a preference for different cell compartments, such as epithelial or immune cells. Some TF classes were unevenly distributed across the cell type specific or ubiquitous TF groups (Fig. f).

In a similar way, TFs can be clustered by shared target genes, so that the identified TF groups regulate overlapping sets of genes. For the top cell-type specific TFs, we find that clusters correspond to cell type compartments where they regulate common target genes as expected (Fig. 1g). For example, combinations of TFs regulate a unique set of target genes across neuronal cell types and TFs involved in eye development cluster together (*OTX2*, *CRX*). Similarly, a group of TFs regulating genes in endothelial cells includes well known endothelial regulators (*SOX17*, *SOX13*), and embeds them in a context with additional co-regulating TFs. The “ubiquitous” TFs found in the largest number of cell types clustered by biological functions (Fig. 1h) such as cell cycle activity (*E2F1*, *TFDP1*, *CTCF*), while others are involved in signalling and transcriptional response, like NFκB and the AP-1 complex, consistent with previous findings^32^. In general, however, “ubiquitous” TFs may be involved in multiple processes and regulate unique target genes, depending on the cell type (Extended Fig. 1b).

Overall, our atlas of cell type and tissue-specific TF regulation programmes highlights the complexity of gene regulation and gives insights into the combinatorial regulation of gene expression, including redundant and co-opted functions of TFs. It allows investigating the broad features of TF regulation from a high-throughput perspective, filter TFs by cell type, and to home in on specific pathways.

### Regulatory map of cell type transitions in the B cell lineage

While TFs can be clustered broadly based on their target genes, the regulons we captured often contain different targets of the same TF for different cell types. This is also true for small transitions from one cell state to the next during development or differentiation, highlighting key regulators at each stage in the differentiation cascade. To illustrate this, we elucidated the TF regulons found during B cell differentiation. Thereby we validate our inferred regulons using the well known biology of B cell regulation, while at the same time embedding known interactions into a wider network of cell-type-specific regulatory factors.

To this end, we selected B cell regulons from CellRegulon, inferred from multiple tissues, including immune-and digestive organs as well as the developing lungs. To group TFs by function, we again used the gene overlap (Jaccard coefficient) as a distance measure between regulons and constructed a nearest neighbour graph for clustering. Clusters therefore represent sets of co-regulated genes linked to groups of TFs which govern the expression regulation of the gene sets.

First, we focused on TFs whose activity changes over B cell differentiation. Unsupervised Jaccard-distance based embedding of regulons arranges them in the correct order of cell type transitions known from previous work, starting from pre-pro-B cells and ending with immature and mature B cells (Fig. 2a). The regulon clusters along this trajectory represent different cell types and TF proportions corresponding to the stages of differentiation. Overall, we identify a network of clusters in which many TFs are co-opted across several cell types^10^ (Fig. 2b). While the same TF may be active in several stages of differentiation, the corresponding number of target genes varies and can be regarded as a measure of importance (degree centrality) of that TF in a given cell type^33,34^. Interestingly, target gene numbers per TF varied along the trajectory in two patterns, consistent with the alternating phases of differentiation and proliferation during B cell development^35^ (Fig. 2b, bottom).

**Figure 2.**
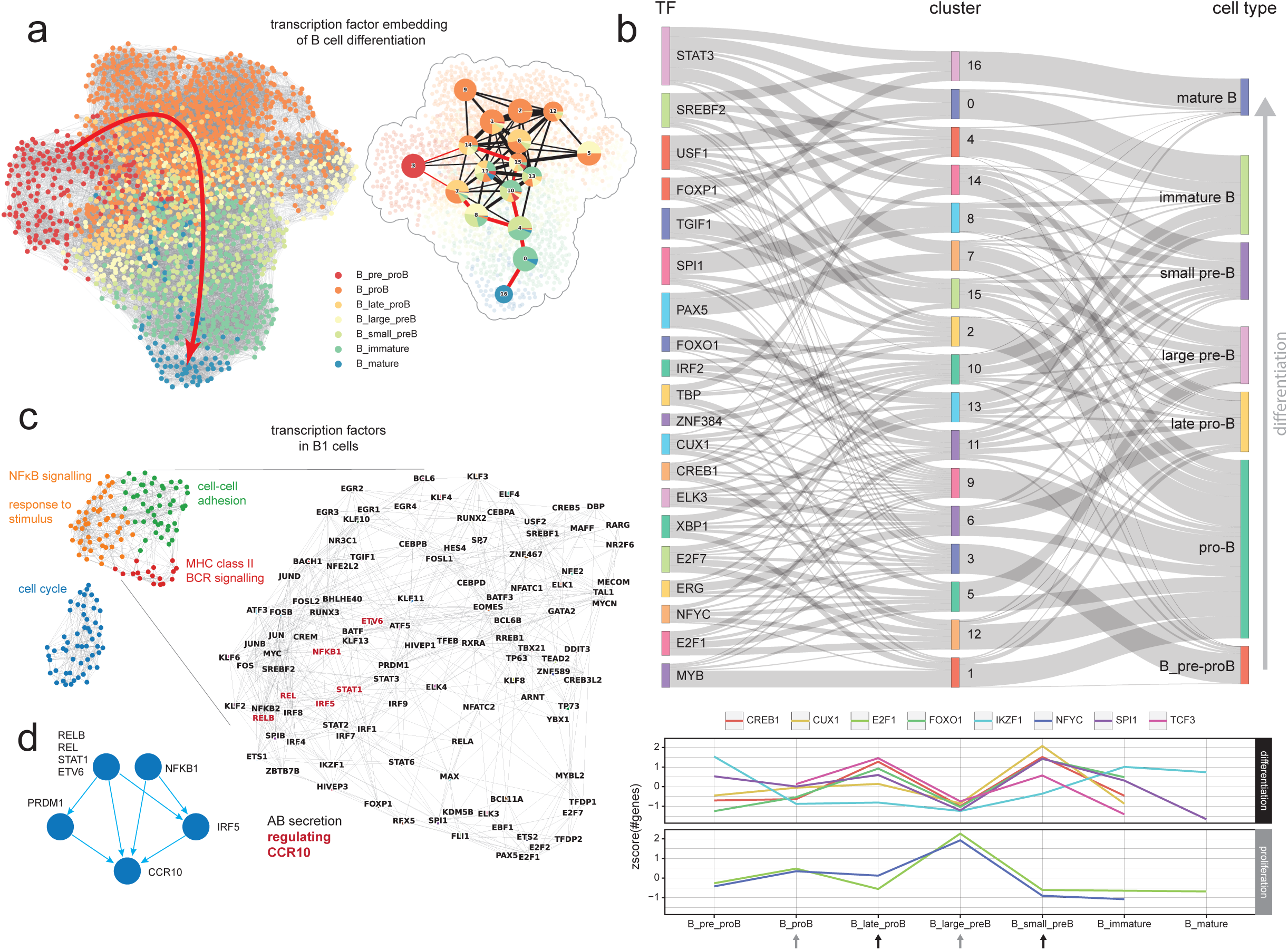
B cell development. **(A)** Network of regulons for B cells. The regulons are arranged using UMAP with Jaccard coefficients of target gene overlap as a similarity metric. By similarity, regulons align in an order that resembles stages of B cell development. Right: clustering of regulons and PAGA graph. Clustered regulons represent a mix of cell types and TFs. **(B)** Sankey-diagram showing the association of clusters to cell types and top TFs. Bottom: variation in the number of target genes along the trajectory for selected TFs. Patterns of variation fall into groups associated with differentiation and proliferation processes that alternate during B cell development. **(C)** Regulon network for B1 cells (arrangement analogous to (A)). Clusters of regulons are annotated for biological processes and TF names are shown for the main cluster. TFs with *CCR10* as a target gene are marked in red. **(D)** Network of TFs regulating *CCR10*.

The TFs identified include classic B cell lineage factors (e.g. *PAX5*, *SPI1*, *FOXO1*, *EBF1*, *IKZF1* and *TCF3*)^36–38^, cell cycle regulators (e.g. E2F family), as well as TFs known to be associated with different stages of B cell differentiation such as *MYB*, *ELK3* and *FOXP1*, driving early commitment, pro-B cell and mature B cell survival, respectively. In addition, we found TFs less commonly associated with the B lineage, such as *CUX1* or *ZNF384*, but which have been reported as biomarkers and fusion partners in B-ALL, in line with their activity in B cell progenitors. Interestingly, key regulators also include TFs that can integrate other stimuli that may affect immune responses such as cAMP signalling (*CREB*)^39^, sterol responses (*SREBF2*) or TGF-beta signalling (*TGIF1*)^40^.

While a large number of studies has investigated the TFs involved in classical B cell differentiation, comparatively less is known about B1 cells which constitute an innate-like type of B cells. Therefore, we aimed to characterise the TFs involved in B1 cell regulation using our atlas. The regulons of B1 cells form two separate clusters, of which one is enriched for cell cycle related genes and one for B cell related functions, including subclusters enriched for different signalling pathways (Fig. 2c). For example, CCR10, a gene recently described as a marker of a B1 subtype with increased spontaneous antibody secretion capacity^22^, was found to be regulated by TFs in a subcluster enriched for NFκB signalling (Fig. 2d).

Overall, this application shows how CellRegulon helps to embed known regulators of B cell biology into a broader context, highlighting the graded, multifactorial and pleiotropic nature of TF regulation. CellRegulon contains a vast number of regulatory interactions from other cell types besides B cells across diverse tissues. The embedding networks can be explored as a resource by cell type, transcription factor, target gene, tissue or dataset, even revealing TFs involved in subtle cell state transitions.

### CellRegulon reveals disease-relevant cell types

Our regulon atlas can also be applied to disease gene signatures to highlight disease relevant cell types and their TFs. Here, we downloaded a published consensus gene signature of adult asthma from nasal and bronchial brushings, containing 1273 differentially expressed (DE) genes^41^ and used it to score the overlap with each regulon in our atlas (Fig. 3a). Since each regulon is annotated with a TF, cell type and tissue, the scores can be summarised for each of these categories.

**Figure 3.**
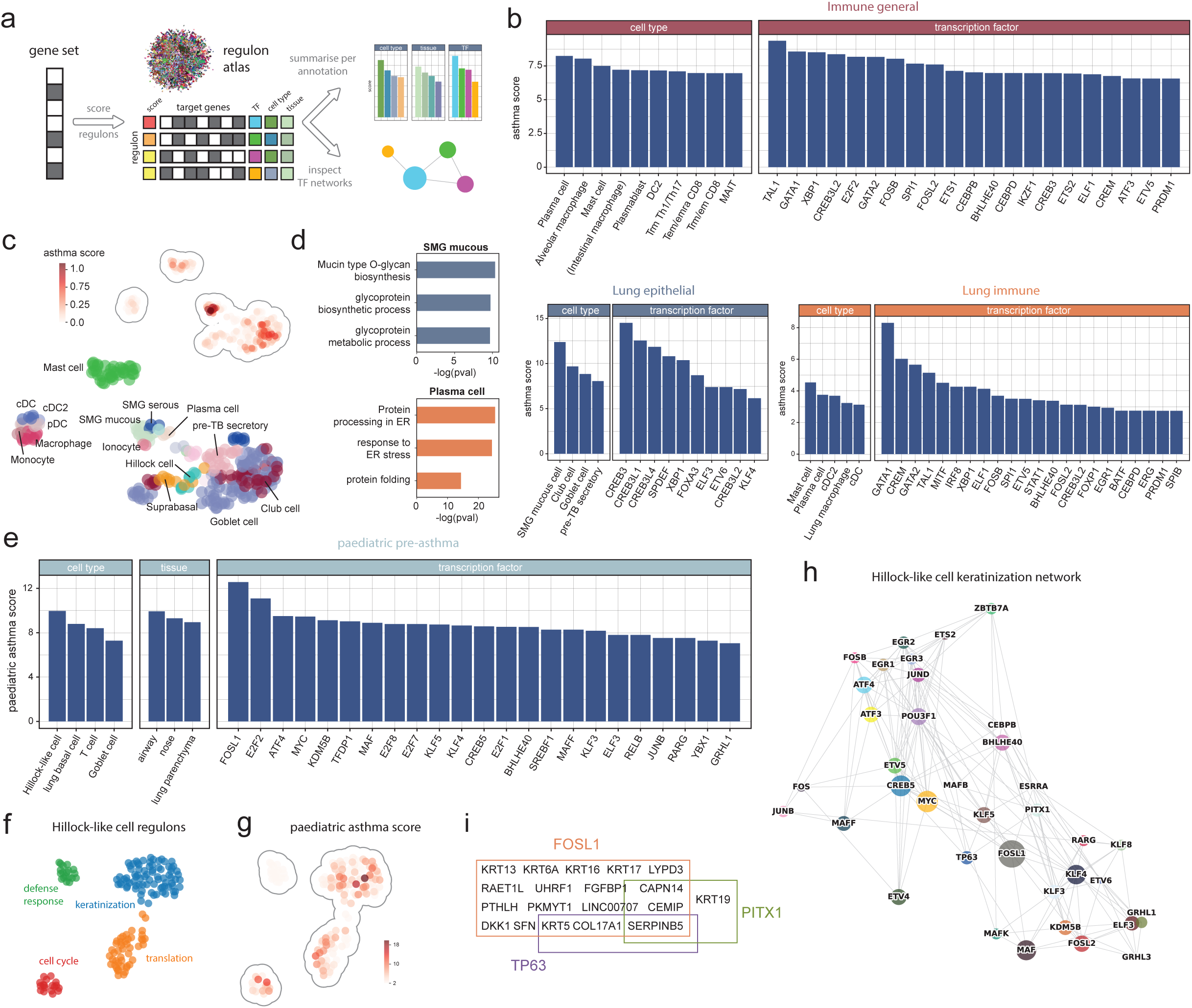
Asthma gene set association. **(A)** External gene sets are scored against the regulon atlas to identify associated regulons based on matching target genes. The TF, cell type and tissue annotations of regulons can be ranked based on the score and visualised in regulon networks. **(B)** Ranking of regulons for a consensus asthma gene set. Top: querying against all immune regulons from the regulon atlas highlights various types of immune cells including plasma cells, macrophages, mast and T cells, as well as associated TFs. Bottom left: querying against lung epithelial cells reveals different types of mucous secreting cells and corresponding transcription factors. Bottom right: querying against lung immune cells highlights different types including mast cells, monocytes and granulocytes, with a highest score for lung megakaryocytes. **(C)** Regulon embedding based on target gene overlap, coloured by cell type. Inset: asthma gene set scores. **(D)** Regulon network for lung megakaryocytes. The node size is scaled by asthma score. **(E)** Ranking of regulons for a childhood-onset pre-asthma gene set. **(F)** Regulon network for Hillock-like lung epithelial cells. Clusters are annotated with biological processes. **(G)** Gene set scores for (F). The highest scores are found in a regulon cluster regulating keratinisation genes. **(H)** Regulon network of the keratinisation cluster. Node size is scaled by gene set score. **(i)** Co-regulated genes for enriched transcription factors.

Summarising by cell type and tissue we find the highest overlap with regulons from immune cells, indicating a relative increase in immune cell numbers in the airway mucosa as expected for adult onset asthma (Fig. 3b top)^42^. Homing into lung epithelial cells, the scores are largest for secretory epithelial cells (Fig. 3b bottom, Fig. 3c). While goblet cell hyperplasia is particularly characteristic in asthma, high scores are found for multiple secretory cells driven by similar TF programs. For example, we find TFs associated with mucus secretion in lung epithelium such as *SPDEF* and *CREB3L1*^43,44^ and target genes including *MUC5AC* and *MUC5B*. Association of regulons from a range of cell types might reflect metaplasia and an altered state of goblet cells that resembles programs in other cell types under these conditions. Interestingly, we find slightly different functions regulated by this TF program across secretory cell types, when looking at their unique target genes. This includes metabolism of mucin production (Fig. 3d) for submucosal gland cells, transport and cell adhesion in goblet cells, as well as epithelium development in club cells (Extended Fig. 3c). In the lung immune compartment, mast, plasma and dendritic cells, as well as lung resident macrophages score especially high, consistent with the literature (Fig. 3b,c)^45^.

Corresponding TFs known to regulate immune functions including GATA1/2, *TAL1*, *IRF8* and *XBP1* rank at the top. For example, *XBP1* shows the highest score for plasma cells, and indeed is a key regulator of plasma cell differentiation, ER expansion and antibody secretion^46,47^, also reflected by its regulon target gene functions (Fig. 3d). In addition, *XBP1* and *SPI1* showed activities during B cell development (Fig. 2b), although their activity signature in plasma cells is more pronounced in this disease state (Fig. 3d).

Interestingly, when including foetal lung regulons, megakaryocyte specific regulations are predicted with a large score (Extended Fig. 3a,b). In mice, lung megakaryocytes have been shown to have different characteristics from bone marrow megakaryocytes including antigen presentation and immunomodulatory activity^48,49^, and have further been suggested to play a role in asthma^50,51^. Focusing on the corresponding TF-network, the highest asthma scores are found in a cluster of TFs including *MAX*, *FLI1* and *ELF1* (Extended Fig 3d). We also note that platelets and their progenitor cells, megakaryocytes, constitutively express IL-33 protein^52^, a mediator of emergency megakaryopoiesis^53^ and central player in asthma^54^, which may explain this observed link. While capturing megakaryocytes close to the epithelial layer under healthy conditions is not expected, epithelial barrier breaches and vascular leakage during inflammatory airway remodeling could bring them closer to the surface, allowing them to exert antigen presenting activity^49^. Future studies will have to validate the functional relevance of these findings.

Next, we investigated scores obtained from a bulk gene signature from pre-asthmatic wheezing children to explore gene regulation at the onset of asthma (Fig. 3e)^55^. This smaller signature of around 400 genes implicated in childhood onset asthma scored highest in regulons from respiratory airway and epithelial cell types. Interestingly, in comparison to adult asthma, the most enriched cell types include basal cells as well as hillock-like cells, a rare cell type that has been linked to proliferation and a squamous differentiation trajectory^56–58^. Consistently, the most enriched TF, *FOSL1 (FRA1)*, is specific for suprabasal and squamous epithelial cells (v23.proteinatlas.org)^59^ and may play a role in inflammation, proliferation and keratinization of these cells^60,61^.

The embedding of hillock-like cell regulons shows four distinct clusters that can be associated with different cellular functions including cell cycle, defence response, protein synthesis and keratinization (Fig. 3f). Among those clusters, the highest disease signature scores are found within the “epidermis development and keratinization” cluster, further supporting a link to squamous differentiation (Fig. 3g). The high-scoring TFs form a network, co-regulating various genes (Fig. 3h), including a large number of keratins such as *KRT13* and *KRT6A*, which are also markers of hillock-like cells (Fig. 3i). Taken together, matching the DE gene signature of childhood onset pre-asthma against our atlas reveals hillock-like cells, as well as *AP-1/FRA1* complex targets, and points to specific dysregulated cellular processes linked to keratinization, in line with recent findings^55^, capturing the distinct origins of adult and childhood onset asthma.

### Extension into a multi-omic regulon atlas of adult lung

While the TF-gene links in CellRegulon already give insights into cell-type-specific gene regulation, additional layers of information can be added to the networks. For example, regions of accessible chromatin can be easily measured using sATAC-seq^62^. Scenic, the pipeline we use for regulon predictions^2^, already considers both correlations in gene expression and the presence of TF binding motifs in the promoters of potential target genes. However, by including single cell ATAC-seq data to define regions of open chromatin, the Scenic+ pipeline allows to include individual enhancer-containing regions in the gene regulatory network^16^. Including accessible regions in enhancer-GRNs (eGRNs) can for example help to evaluate the effect of genetic variation in non-coding regions of the genome.

To investigate regulatory interactions and their role in lung disease further, we subsetted lung regulons from CellRegulon and extended them into an eGRN. To this end, we created a new single cell atlas of the adult lungs profiled with 10X multiome (RNA & ATAC). We then made use of public GWAS summary statistics to calculate disease risk from open chromatin, and propagated these signals over the gene regulatory network to identify disease-relevant sub-networks.

To create a multi-omic lung atlas, we processed samples from 5 adult donors across 5 locations using 10x multiome, capturing both RNA and ATAC signals from the same cell. After QC filtering this new multi-modal atlas of the adult lungs comprises over 37,000 high quality cells. Utilising several published datasets as a reference^25,63,64^, we annotated 60 cell types, including large numbers of epithelial, immune and endothelial cells as well as fibroblasts specific to the lungs (Fig. 4a). The annotation also includes rare and recently discovered cell types such as hillock-like cells^56–58^, AT0^65^ cells and distal secretory cells^66^. We then employed the SCENIC+ pipeline^16^ to predict gene regulatory links comprising genes and accessible genomic regions, making use of both the RNA and ATAC modalities. The multiomic single cell atlas of the adult lungs and eGRN are made available as resources particularly suited for the exploration of multimodal gene regulation across locations of the adult lung.

**Figure 4.**
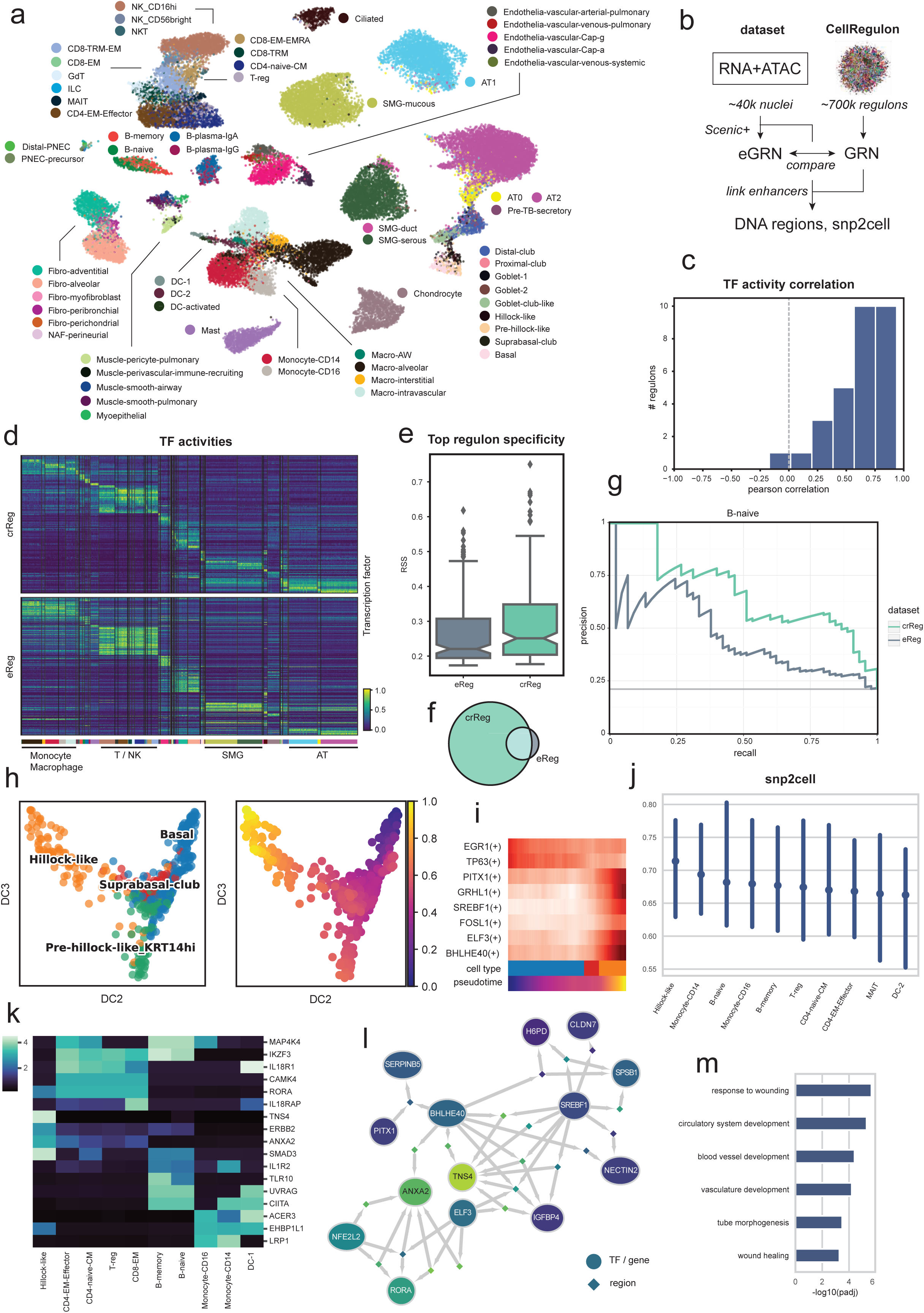
Analysis of eGRNs using multiome data. **(A)** UMAP of the adult lung multiome atlas showing cell type annotations. **(B)** Flow diagram illustrating analysis steps and comparison between multiome atlas enhancer-regulons and CellRegulonDB. **(C)** Distribution of correlations between TF activities over the lung atlas using eReg and crReg regulons for the 50 most highly variable TFs. **(D)** eReg and crReg TF activities over the lung atlas. CellRegulon regulons differentiate cell types with a finer resolution. **(E)** Boxplot of regulon specificity scores (RSS) for the top 3 TFs per cell type, showing higher specificity for crReg regulons. **(F)** Venn diagram showing the overlap of cell type specific TFs for eReg and crReg regulons with RSS>0.25. **(G)** Precision-recall curve evaluating prediction of TFs relevant in naive B cells using text mining (ChatGPT). **(H)** Diffusion map of the trajectory for hillock-like cells. Left: cell type annotation. Right: diffusion pseudotime. **(I)** TF activities along the axis between Basal and hillock-like cells. **(J)** Distributions snp2cell scores per cell type, ordered by median (dots). **(K)** Heatmap of snp2cell scores per gene per cell type. **(L)** eGRN showing a sub-network of TFs enriched for COA GWAS scores. **(M)** Gene set enrichment (GO BP) of the top 10 genes with largest COA GWAS score.

In addition to disease-risk analysis, we first compared the CellRegulon network with the gene regulatory network inferred from only the new dataset (Fig. 4b). Inferring a GRN from only paired RNA and ATAC data should help to improve TF-target-gene predictions. On the other hand, before adding the ATAC data, CellRegulon was derived from a much larger collection of RNA-based datasets comprising over 100x more cells and therefore likely captures a larger variability of gene expression states. To compare the two GRNs, we first computed the correlation between TF activities obtained using regulons from the multiomic dataset (eReg) and the CellRegulon atlas (crReg) across all cells in our multiomic lung atlas for the most highly variable TFs. We generally find large positive correlations for most of the TFs (Fig. 4c). Visualising the activities of the top TFs per cell type (Fig. 4d) we find similar patterns with blocks of related cell types that are difficult to distinguish based on TF activity, such as different T-cell subsets. Interestingly, the crReg regulons show a finer distinction between some of the cell types, which is further reflected in significantly larger regulon specificity scores (RSS) when selecting all top 5 TFs per cell type (Fig. 4e, Wilcoxon rank-sum test, p-value: 0.0036). Therefore, when nominating TFs per cell type based on high specificity, around 7x more TFs can be selected from CellRegulon (Fig. 4f). This shows the advantage of using a larger dataset collection for regulon inference and also reflects the availability of a larger pool of cell type specific regulons. To further address the question, whether this TF selection indeed reflects biology, we compared predictions against the biological literature. Using B cells as an example that has been extensively studied in the past, we created a large ground truth set of 64 TFs with a role in naive B cells using semi-manual text mining (see Methods). The comparison between crReg and eReg showed an advantage of crReg, with a larger precision at the same recall values (Fig. 4g), suggesting that more relevant TFs can be recovered.

These results indicate that an increased data volume is beneficial for recovering regulatory interactions more completely. A challenge in computational GRN inference is the low overlap of identified connections, both between methods and for different runs of the same method^67^, which is aggravated by the immense size of the universe of possible gene interactions. As the expression variability in each dataset likely only reflects a small portion of this universe, inferring regulation from analyses of multiple datasets and broken down into cell lineages, as done for CellRegulon, may therefore help to arrive at a more comprehensive and robust sampling of the full regulatory landscape.

We next investigated individual cell states in our multiomic lung cell atlas further using regulons from CellRegulon. To this end we subset cells that lie on a keratinization trajectory from lung basal cells to hillock-like cells (Fig. 4h). Basal cells, suprabasal cells and hillock-like cells form a continuous transition in diffusion space. To examine TF activities along this continuum we defined pseudotime from basal to hillock-like cells (Fig. 4i), although recent findings suggest that the reverse transition is also possible^58^. TFs showing higher activities at the start of the trajectory including *TP63* and *EGR1* are linked to basal cell maintenance and suppression of inflammatory responses^68,69^, while TFs with roles in keratinization, such as *FOSL1*, *SREBF1* or *BHLHE40*^59,60,70^, are more active towards the end of the trajectory.

Another use case for CellRegulon is the GRN-aware mapping of genetic risk data from GWAS to specific cell types. To further investigate links between hillock-like cells with childhood-onset asthma using published GWAS summary statistics^71^, we transformed the crReg regulons into an eGRN by adding peak-gene links from co-accessibility across our multiomic lung atlas (Fig. 4b). The resulting enhancer-GRN allows to identify TFs targeting many genomic regions with a large disease association. To this end, we used snp2cell^72^, a method that identifies cell type specific disease-associated TFs. Interestingly, hillock-like cells show the largest (Fig. 4j) median association across all genes, suggesting that many disease-relevant pathways are active in this cell type.

In this analysis, the top enriched TFs and genes cluster into cell type-specific groups (Fig. 4k). Genes implicated in asthma that regulate immune related functions like *IL18R1*^73^, IL1R2 (which can bind IL33^74^), *IKZF3*^75^ and *TLR10*^76^ are highlighted for T cells, monocytes and other immune cells. In hillock-like cells, a distinct set of genes is enriched including *ERBB2*, *ANXA2*, *TNS4* and *SMAD3*. Epithelial cells from asthmatics were shown to exhibit impaired wound repair, linked to *ERBB2*^77^. *ANXA2* is linked to Ca^2+^ signalling, inflammation and tissue repair^78^. Relatively little is known about *TNS4*, however, this focal-adhesion-linked gene binds keratins and was shown to promote keratinocyte proliferation^79^. Methylation in the promoter of *SMAD3* has been linked to asthma in children after rhinovirus-induced wheezing^80^ and, as part of TGFβ-signalling, regulates inflammation in wound repair through inhibition of AP-1 activity^81^. Visualising the top enriched TFs in hillock-like cells in a network (Fig. 4l) shows connections between many of the genes and additional disease associated TFs with increased activity towards the end of the keratinization trajectory (Fig. 4i), including *ELF3*, *BHLHE40*, *PITX1* and *SREBF1*. These results further support a role of the keratinization trajectory and hillock-like cell state in the early stages of asthma. Gene set enrichment of GO biological processes^82,83^ for the hillock-like cell specific asthma associated genes further shows wound response among the top terms (Fig. 4m). This could suggest the keratinization trajectory is triggered by an injury to the lung epithelium, such as a viral infection. Examining the meta-data of our lung cell atlas, we find increased expression of hillock-like cell marker genes in donors with a positive smoking-status (Extended Fig.4), which is confirmed when looking at a larger integrated lung cell atlas (HLCA v1)^25^. Thus, injuries such as through smoking or possibly environmental pollutants might trigger a wound response and keratinization^84^, and in some cases increased inflammation leading to the chronic changes associated with the development of asthma.

### Gene regulatory networks in lung disease

Risk for complex diseases such as asthma, COPD and IPF is influenced by a large number of inherited genetic variations, or single nucleotide polymorphisms (SNPs), potentially affecting biological processes across many cell types. Such genetic variations are mostly present in non-coding regions of the genome, including enhancers. To examine aggregated disease associations over TFs, genes and regions in our gene regulatory networks, we made use of publicly available GWAS summary statistics for COPD, IPF and asthma.

Signals from individual genomic locations can be aggregated over a network using network propagation to increase robustness of the results^85^. We used snp2cell, which performs propagation of risk scores over a gene regulatory network to find sub-networks with overlapping GWAS and cell type enrichment scores^72^. This approach allows to predict “risk” TFs that may not be affected by genetic variations themselves, but that regulate a large number of genes associated with risk SNPs, implying that these TFs are relevant for the studied disease. In addition, cell-type-specific TFs and genes with a disease association can be identified as well.

Comparing asthma, IPF and COPD disease scores, derived using CellRegulon regulons together with our newly generated ATAC data, highlights cell type specific activity patterns (Fig. 5a, Extended Fig 5a-c) indicating which cell populations may be affected. IPF associated genes show strong activity patterns in secretory cell types, including goblet and club cells, as well as submucosal gland (SMG) mucous cells, mainly due to GWAS associations for secreted proteins such as *MUC5B* and *MUC5AC*. Indeed, *MUC5B* promoter SNPs are the strongest genetic risk factor for IPF^86^*. MUC5B* and other mucins including *MUC5AC* have been implicated in impaired mucociliary clearance as a key pathogenetic factor^87,88^. Additionally, IPF scores are high across different types of T cells^89^ and alveolar macrophages^90^. However, as IPF is a rare disease, the size of GWAS studies has been comparatively small and the power to detect all relevant associations may be limited.

**Figure 5.**
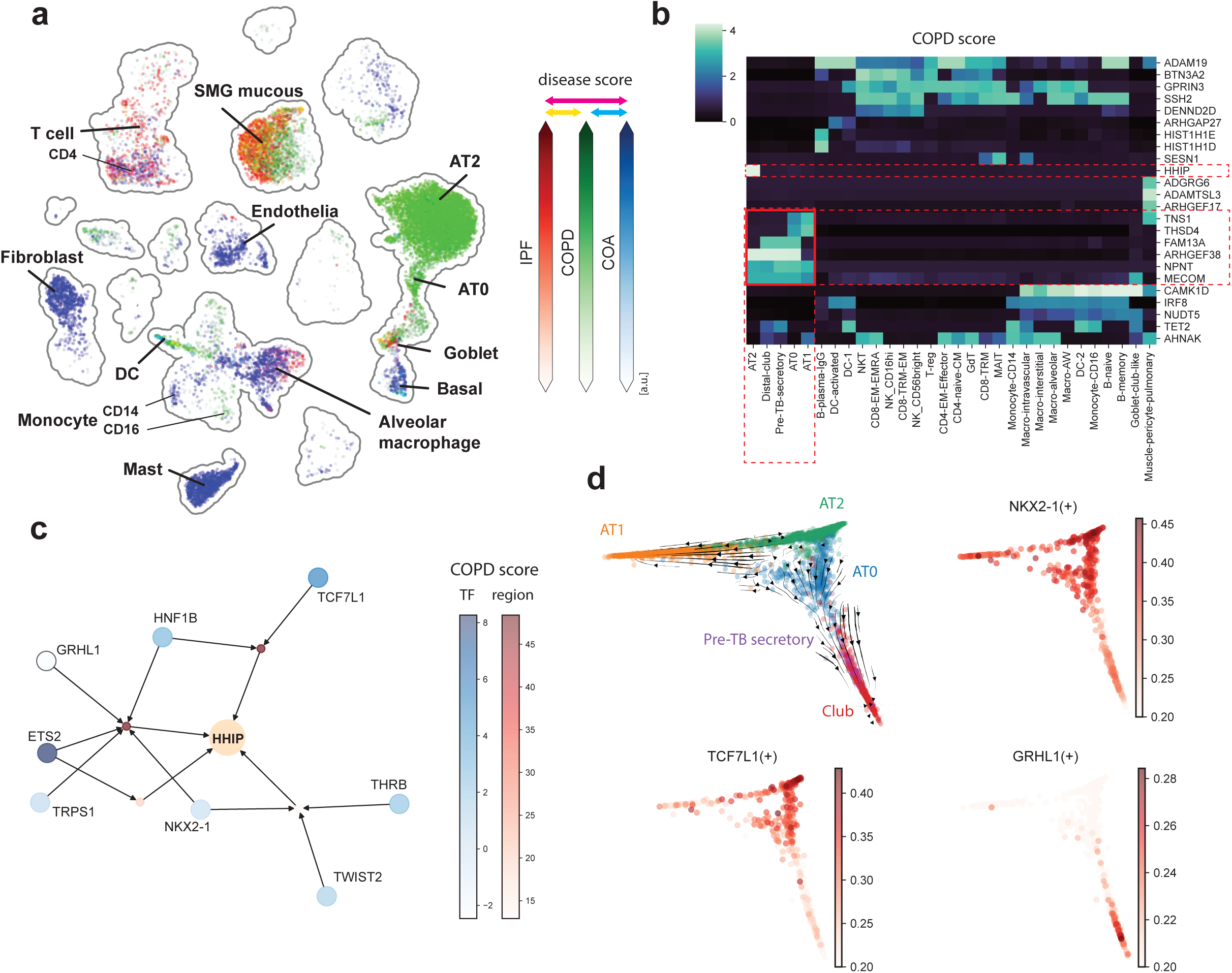
Genetic TF-associations in lung disease. **(A)** UMAP of the adult lung multiome atlas showing GWAS scores for IPF, COPD and COA inferred with snp2cell. The three diseases are shown in different colors, with overlaps shown using additive color mixing. **(B)** Heatmap of COPD scores for genes per cell type. A cluster of alveolar type and secretory cells is highlighted in red. **(C)** GRN centered around the HHIP gene. TFs and genomic regulatory regions with predicted TF-binding are shown as large and small circles respectively. TFs and regions are colored by snp2cell COPD score. Note, that TFs usually have smaller scores than regions, since they participate in a multitude of interactions. **(D)** Diffusion maps of the alveolar type differentiation trajectory surrounding AT0 cells. Top left: scVelo streamline plot showing directions inferred by splicing dynamics. Others: activities of selected TFs inferred by AUCell.

For COPD, disease scores overlapped with IPF on secretory cells (goblet, SMG mucous) and were highest in alveolar type 2 (AT2) cells, respectively pointing to chronic bronchitis and emphysema phenotypes^95^. In addition, various immune cells including B cells, dendritic cells and CD16+ monocytes score high, as expected. Alveolar fibroblast scores are slightly elevated, but not high compared to other cell types, indicating that characteristic changes of fibroblasts observed in COPD may be predominantly signalling-induced and epi-genetic^91^, rather than a consequence of germline variation. For childhood-onset asthma (COA), risk scores for epithelial cells were highest in basal cells and also in hillock cells (Fig. 5a, Extended Fig. 5b), consistent with our previous findings (compare Fig. 4). Amongst the immune cells, scores are high in B and T cells, as expected^92^, and signals overlap with IPF for alveolar macrophages. In contrast to IPF and COPD, particularly large asthma scores are found in mast cells, pointing to the allergic component of the disease^93^. Overall, snp2cell score distributions recapitulate known disease-cell-type associations^94^, providing a large-scale overview over characteristic differences and commonalities.

In addition to cell types, TFs and genes within the enriched CellRegulon networks can be explored. In the epithelial compartment, all three diseases show large scores with different distributions (Fig. 5a). A distinct set of disease associated genes overlaps with AT2 specific expression profiles (Fig. 5b), including *FAM13A* and *HHIP*, well known COPD genes with roles in epithelial repair through WNT/β-catenin and TGF-β signalling^96,97^, whose function in COPD pathogenesis is still under active research. *MECOM* was also enriched and has been linked to lung function in GWAS^98^, although its role in COPD pathogenesis is unknown.

However, *MECOM* (*PRDM3*, *EVI1*/*MDS1*) has been shown to interact with TGF-β^99^ and WNT/β-catenin signalling in cartilage^100^, and to regulate *NKX2-1* mediated alveolar epithelial differentiation^101^. Overall, disease scores in this cluster point to a role of impaired alveolar regeneration in COPD via WNT/β-catenin and TGF-β signalling involving various transcription factors.

In addition to linking TFs with COPD, snp2cell also calculates enrichments for genomic locations in the GRN with a predicted regulatory function. Focusing on the regulatory interactions around *HHIP* (Fig. 5c), a well known COPD-associated gene, this prioritises two genomic locations bound by *HNF1B*, *TCF7L1*, *ETS2* and *NKX2-1* among others. *NKX2-1* is known as a master regulator of AT2 cells that helps to maintain cell identity and stemness^102^. *TCF7L1*, a component of the WNT pathway, is dysregulated in COPD^103,104^. The embedding of COPD signals in a sub-GRN related to epithelial cell differentiation of alveolar type cells further highlights the role of impaired alveolar repair in emphysema type COPD and puts the regulation of individual genes and regions into context.

Interestingly, a cluster of alveolar epithelial cells and distal secretory cells shares multiple enriched genes (*TNS1*, *THSD4*, *FAM13A*, *ARHGEF38*, *NPNT*, *MECOM*; Fig. 5b). While some of the genes are specific to AT2, AT1 or secretory cells, AT0 cells have the highest enrichment of these genes. AT0 cells have been described as intermediate states on a trajectory from AT2 to AT1, and secretory cells^65^ or vice versa^66^. While both transitions may be possible, splicing dynamics in our healthy multi-omic atlas suggest a transition away from AT2 cells (Fig. 5d). TF activities of *NKX2-1* and *TCF7L1* with GWAS enrichment are also highest in AT2 and AT0 cells (Fig. 5d), presumably contributing to maintenance of stemness. In contrast, GRHL1, was also found to bind to this regulatory region, but was more active in distal secretory cells and also showed a lower COPD association score. Thus, together with GWAS scores, the distribution of TF activities over the trajectory of alveolar differentiation further illustrates how decreased stemness of AT2 cells might lead to impaired alveolar repair under COPD conditions thereby promoting emphysema development.

### CellRegulonDB: an interactive online resource of human cell type specific regulons

The regulon atlas analysed in this work, which has been created using a large collection of human cell atlas data, has been made available for easy access and interactive browsing through a web-interface (https://www.cellregulondb.org). The web-interface allows querying for specific transcription factors, target genes, tissues and cell types, and displays subsets of the network, which can be downloaded as a table or PDF. Furthermore, CellRegulon can be accessed and analysed programmatically through a corresponding python package (https://github.com/Teichlab/cellregulondb) for more complex analyses. Regulons can be downloaded and formatted using the python package to facilitate various downstream analyses. The package also provides basic functions for analysing the data, including looking up TFs and their cell type-and tissue-specific target genes, ranking and clustering and finding enriched biological functions. Finally, matching regulons with custom gene signatures, such as DE-based disease gene sets can be performed through the python package, and networks can be exported for GWAS-based risk analysis with external tools^72^.

## Discussion

Transcription factors play a crucial role in shaping cell states and directing differentiation. Many regulate their target genes in a cell type specific manner and in varying combinations with other TFs. To resolve this complexity, experimental methods have been employed, including cell type specific ChIP-seq experiments, CUT&RUN^19^ or TF overexpression screens in iPSCs. However, the combinatorial nature and cell state dependence make it difficult to test TF effects across all conditions in the lab. Computational methods present another opportunity, using large-scale datasets to obtain predictions of TF regulation.

Increasingly large amounts of single cell data produced through initiatives such as the Human Cell Atlas make it possible to cover an extensive number of cell types with this approach.

Here, we present an atlas of TF regulation computed across 15 organ systems and over 700k cell type-and tissue-specific transcription factor-target gene sets. The atlas is available as an interactive resource for the research community. While computing GRNs *de novo* often requires large amounts of computational resources, the pre-computed regulons can be accessed and investigated on a standard computer in minutes. Using query data, matching regulons can be extracted across a large number of cell types. Overall, our paper makes the following contributions: (i) We describe CellRegulon as a resource; (ii) We characterise the overall distribution of TFs across cell types and classify them into specific and ubiquitous TFs based on their occurrence patterns; (iii) We demonstrate how to use the regulon atlas on its own, in combination with external gene sets or datasets to obtain predictions of TF regulation, for example for understanding asthma relevant TFs and cell types; and (iv) we created a new multi omic single cell atlas (RNA+ATAC) of the adult lung, which we used to extend the precomputed GRNs for lung cell types to a set of lung enhancer-GRNs. These GRNs were used to analyse gene regulation along a trajectory of epithelial keratinization, a process we show to be enriched with a childhood-onset asthma score. (v) Furthermore, lung eGRNs were used to compute disease risk per cell type for childhood-onset asthma, IPF and COPD, predicting specific risk gene modules within these eGRNs.

The atlas of cell type specific regulons reveals a dynamic landscape of tight cross-regulation in which many TFs are co-opted across cell types, with often varying sets of target genes and in varying combinations with other TFs. However, most TFs are predicted to be specific for only a small number of cell types and to regulate a smaller number of genes compared to ubiquitous TFs. This split reflects either more cell type specific functions, such as eye photoreceptor development (VSX1), or universal ones, such as the cell cycle (E2F1). While this principle has been described before^17^, our atlas constitutes a high-resolution map that allows zooming-in and investigating specific cell types and TF-combinations in detail.

With over 700k regulons across more than 500 human cell types it spans a vast space to explore, in which we focused on a few selected examples. We show that regulons recapitulate the trajectory of B cell maturation and comprise TFs that are well studied in this context, thereby validating our approach. These are embedded in a larger network of TFs, suggesting potential interactions and regulatory relationships. Further, we screen regulons of B1 cells to find TFs that may regulate cell type markers linked to increased antibody secretion. Finally, we filter out regulons enriched for disease gene sets of asthma, revealing immune and secretory epithelial cell types with their corresponding TFs. For pre-asthma we identify TFs linked to lung epithelial keratinization and hillock-like cells which we further investigated using our multi-omic lung cell atlas. Combining ATAC data, gene regulatory networks and GWAS data of early asthma we highlight a regulatory sub-network linked to keratinization and wound response, suggesting a role of epithelial injury in combination with a hereditary component in asthma-inception. Like our regulon atlas, the multi-omic lung enhancer-GRN constitutes a resource for further exploration of regulatory relationships.

These resources we provide here allow the assessment of regulatory mechanisms across many cell types, genes and TFs, but importantly, suggested mechanisms remain predictions and require experimental validation. The large-scale nature of these resources makes them particularly suitable for hypothesis generation, making predictions or selecting candidates for further validation. CellRegulon allows linking single cell atlases with disease or perturbation gene signatures as well as GWAS summary statistics, generating new insights via a pre-computed resource. It also allows extracting subsets of TFs and genes for more detailed mathematical modelling. A limitation of the CellRegulon approach is that some quantitative details of gene regulation, such as the relative contribution and cooperativity between TFs, kinetic parameters and target degradation are not captured in the regulatory graphs.

However, CellRegulon can be used to survey and home in on subsets of TFs and genes for more detailed mathematical modelling.

Going forwards, gene regulatory network prediction is likely to involve more advanced machine-learning methods, including artificial neural network models^105–107^, which are a current area of active research. Here, we utilized the pyScenic pipeline^2^, an established machine-learning method for gene regulatory network inference based on gradient boosting machines, which allow easy extraction of feature importances. As more and larger datasets become available for model fitting, we expect that more complex machine learning models will become more widely used. These may contribute to future versions of the CellRegulon database. Conversely, GRNs from CellRegulon may also serve as prior knowledge for machine learning approaches, such as graph neural networks or with frameworks similar to CellOracle^108^.

CellRegulonDB provides a snapshot of the gene regulatory landscape of the whole body, across far more tissues and with much greater cell-type-specificity than previous efforts. We predict that resources like the regulon atlas, together with advancements in machine learning and artificial intelligence, will increase our insights from single cell data and contribute to many applications in basic research, drug development and tissue engineering.

## Acknowledgements

This project has received funding from the European Union’s Horizon 2020 research and innovation programme under the Marie-Skłodowska-Curie Action grant agreement No. 101026233 to J.P.P. and grant agreement no. 874656 (discovAIR) to K.B.M and M.C.N. H.Y. acknowledges funding from the European Union’s Horizon 2020 research and innovation programme under grant agreement no. 859860 (CANCERPREV). M.K. acknowledges funding from the LEO Foundation (LF-OC-19-000225), Swedish Research Council (2022-01059) and Swedish Cancer Society (21 1821Pj). M.C.N. acknowledges funding from the Ministry of Economic Affairs and Climate Policy (the Netherlands) through a PPP-allowance from the Top Sector Life Sciences & Health, the Netherlands Lung Foundation project 4.1.18.226 and the Chan Zuckerberg Initiative, LLC Seed Network grant CZF2019-002438 “Lung Cell Atlas 1.0”. K.B.M. and S.A.T. were supported by the Wellcome Trust (WT211276/Z/18/Z and Sanger core grant WT206194). S.A.T. received funding from the Chan Zuckerberg Foundation (2019-002445, 2022-249170), Wellcome Trust (220540/Z/20/A) and the CIFAR Macmillan Multi-scale Human Program.

## Data Availability

High-throughput raw sequencing data in this study are available from ArrayExpress (www.ebi.ac.uk/arrayexpress) with the accession number E-MTAB-14583. Processed multiome RNA and ATAC-seq data are available for visualization and downloaded from CELLxGENE. (Both raw and processed data will be released upon publication).

## Methods

### Access to human tissue and ethics oversight

Samples were obtained from deceased transplant organ donors by the Collaborative Biorepository for Translational Medicine (CBTM) with informed consent from the donor families and approval from the National Research Ethics Services (NRES) Committee of East of England, Cambridge South (15/EE/0152). CBTM operates in accordance with UK Human Tissue Authority guidelines.

### Tissue collection

Tissue was collected from 5 donors from five lung locations including trachea, bronchi at the second/third generation, bronchi at the fourth generation, upper left lobe parenchyma and lower left lobe parenchyma (Table or Supp Figure with tissue overview). Following collection at the clinic, samples (range: 1–4Lcm3) were immediately placed into cold Hypothermasol FRS32. Within 12 h after circulation ceased, samples were preserved in optimal cutting temperature (OCT) compound and frozen in isopentane at −60 °C for later nuclei isolation.

### Multiome single nuclei sequencing

Our single nuclei isolation method from frozen tissue used 16 × 50 μm thick sections which were homogenized using a glass Dounce homogenizer (Sigma) in nuclei isolation buffer (NIM; 0.25 M sucrose, 0.005 M MgCl 2, 0.025 M KCl, 0.01 M Tris (buffer pH7.4), 0.001 M DTT and 0.1% Triton X-100) in the presence of Complete protease inhibitors (Roche) and RNAse inhibitors RNasin (Promega) 0.4 U μl^−1^ and SUPERase-In (Invitrogen) 0.2 U μl^−1^). Tissue was homogenized using ∼15 strokes with pestle A (clearance 0.0028–0.0047 in.) and then pestle B (clearance 0.0008–0.0022 in.). Isolated nuclei were filtered through a 40 μM filter, collected at 500g and resuspended in 0.5 ml of storage buffer (PBS containing 4% BSA and RNasin (Promega) 0.2 U μl^−1^). Nuclei were incubated with NucBlue (ThermoFisher) and purified from debris by FACs sorting, stained with Trypan blue and counted.

Five thousand nuclei from five different samples were pooled and all 25,000 nuclei were further processed for 10x Chromium Next GEM Single Cell Multiome ATAC + Gene Expression according to the manufacturer’s protocol (reference: CG000338). Nuclei suspensions were loaded with a targeted nuclei recovery of 15-16,000 droplets per reaction (to recover ∼3,000 nuclei per sample). Samples were later demultiplexed by genotype.

Quality control of cDNA and final libraries was carried out using Bioanalyzer High Sensitivity DNA Analysis (Agilent). Libraries were sequenced using a NovaSeq 6000 (Illumina) with a minimum sequencing depth of 20,000 read pairs per droplet.

### Multiome data processing

Sequencing data were aligned to the human reference genome (GRCh38-2020-A-2.0.0) using CellRanger-ARC software (v.2.0.0). The called barcodes from 10x multiome lanes with pooled genotypes from samples from multiple sample donors were demultiplexed per Genotype using BAM outputs through Souporcell (v2.0). Subsequently, the Souporcell outputs were clustered by genotype for metadata assignment to each barcode. For gene expression data, SoupX was applied to remove background ambient RNA. For cellranger-arc called matrices that contained >16,000 droplets (exceeding the number expected from targeted droplet recovery) we increased the estimated global rho value by 0.1 to account for the potential of additional ambient RNA. Droplets were filtered for >200 genes, and <20% mitochondrial reads. Doublet removal is described later. For scATAC-seq, we applied ArchR138 (v1.0.2) to process the outputs from Cellranger-atac. Initial per-cell quality control was performed considering the number of unique nuclear fragments, signal-to-background ratio and the fragment size distribution. Moreover, cells with TSSenrichment score<7 and nFrags< 1000 were removed. Doublets were discarded using the default settings. Initial clustering was performed at resolution = 0.2 with top 40 dimensions from Iterative Latent Semantic Indexing (LSI). Then, pseudo-bulk replicates were made for each broad cell type per region from the initial clustering results. Peak calling (501 bp fixed width peaks) was performed based on pseudo-bulk coverages by macs2. Then, a cell-by-peak count matrix was obtained and exported.

### Computation of regulons

To create a regulon atlas that covers an extensive list of tissues and cell types, we collected single cell datasets from 14 organ systems, prioritising already integrated large single cell atlases with harmonised annotations and supplemented them with additional large studies^20–28^. To infer regulons for specific cell types, datasets were divided into lineages or cell compartments and we employed pyScenic^2^, which computes TF-target gene links based on co-expression, as well as enrichment of TF binding sites around the gene promoter. Inference was performed per lineage to account for biological differences, while also striking a balance between granularity and sufficient data volume. Single cells were further grouped into mini-metacells (10-15 cells) to mitigate data sparsity and make this computationally intensive analysis more feasible. The cell compartment specific regulons were then further filtered for active regulatory interactions by subsetting target genes using differential expression (DE) per cell type and tissue. Since the co-expression based inference itself does not rely on cell type labels and instead requires counts for all genes as input, no integration and common embedding for cell type harmonization had to be performed. The regulon database was instead set up as a pool of gene regulatory TF signatures reflecting the variation found in individual contributing datasets processed separately, which may help to overall capture a larger number of regulatory connections probed under slightly different conditions. For exploration, regulons can be aggregated across multiple conditions by filtering target genes based on their frequency, to prioritise robustness or completeness, where frequency 1.0 and 0.0 would correspond to the intersection and union of target genes, respectively. Cell type labels have been harmonized manually by mapping to Cell Ontology IDs^109^, while also retaining the original author annotation. This approach ensures that new datasets can more easily be added and results linked to the individual publications underlying CellRegulon.

### Regulon analysis

For easy exploration and analysis, the regulon atlas was loaded into an AnnData^110^ object, with target genes as “var_names”, regulons as “obs_names” and a sparse binary data matrix, where “1” represents that a target gene is included in the regulon. Additional annotations including TF, cell type and tissue that a regulon belongs to were stored in the “obs” field of the AnnData object. Therefore, the regulon atlas could be analysed with Scanpy^110^ in a similar manner as conventional single cell atlases. A wrapper class and convenience functions to facilitate regulon-related analyses are available as part of our companion package in python (https://github.com/Teichlab/cellregulondb). Regulons were filtered for a minimum of 4 target genes for all analyses, to exclude noisy connections.

Embeddings to represent the similarity of regulons were computed using “sc.tl.neighbors” and “sc.tl.umap” in scanpy, and clusters computed with “sc.tl.leiden”; with the exception that Jaccard coefficients were used as a similarity metric (thus measuring the overlap of regulon target gene sets) instead of euclidean distances. Trajectories over regulons were computed using PAGA^111^ with “sc.tl.paga” on Leiden^112^ clusters. Selecting regulons matching external gene signatures was also done in analogy to single cell data analysis, using “sc.tl.score_genes”. In the case of a binary regulon membership matrix, this method translates into scoring the fraction of target genes for each regulon that are contained in the external gene signature and comparing it to the fraction obtained by random, while adjusting for the frequency of target genes across regulons, so that more unique genes are weighted higher.

Heatmaps were computed in python using seaborn and scipy for clustering and optimal leaf ordering^113^. Circular heatmaps and dendrograms were computed using the circlize package in R^114^. Sankey plots were created in python using plotly (https://plot.ly). Gene set enrichment (overrepresentation) analysis was performed using g:Profiler^115^ (version e112_eg59_p19_25aa4782) with GO:BP^82,83^ and KEGG^116^ gene sets.

### Cell type annotation

Newly generated single cell 10X multiome data was processed using Scanpy^110^. QC was performed, filtering out cells with less than 200 genes or over 20% mitochondrial genes, and genes found in less than 3 cells, and doublets were removed using scrublet^117^. Batch correction was performed in scanpy by computing highly variable genes per batch, regressing out cell cycle effects and applying Harmony. After embedding, an additional round of doublet removal was performed by overclustering cells with Leiden resolution 10, and removing clusters where >30% of cells have a scrublet score above 0.2. After that, cell type annotation was performed by using a combination of automated label transfer with CellTypist^118^ based on three published lung datasets^25,63,64^ and manual annotation based on known marker genes. To this end, the complete dataset was repeatedly sub-clustered and clusters compared with mapped labels and examined for expressed marker genes. In total, 60 cell types have been annotated.

### eRegulon analysis

Adult lung 10X multiome RNA and ATAC data was preprocessed as described above. We then used the Scenic+^16^ (v1.0.0) pipeline to predict transcription factors and putative target genes as well as regulatory genomic regions with TF binding sites. Meta-cells were created by clustering droplets into groups of around 10–15 based on their RNA profiles and aggregating counts and fragments per metacell. CisTopic (pycistopic v1.0.2) was applied to identify region topics and differentially accessible regions from the fragment counts as candidate regions for transcription factor binding. CisTarget (pycistarget v1.0.2) was then run to scan the regions for transcription factor-binding sites, and GRNBoost2 (arboreto v0.1.6)^119^ was used to link transcription factors and regions to target genes based on co-expression or accessibility. Enriched transcription factor motifs in the regions linked to target genes were used to construct transcription factor–region and transcription factor–gene regulons. Finally, regulon activity scores were computed with AUCell based on target gene expression and target region accessibility, and regulon specificity scores (RSS) derived from them within the Scenic+ framework.

### GWAS association analysis

Disease association analysis for TFs, genes, regions and cell types based on GWAS summary statistics was performed using snp2cell^72^. A Bayesian approach to summation of GWAS signals per genomic region was applied as described previously^120^ and implemented in the nf-fgwas nextflow pipeline (https://github.com/cellgeni/nf-fgwas). Published GWAS summary statistics were obtained for childhood-onset asthma (COA)^71^, IPF^121^ and COPD^122^ (EBI GWAS catalog IDs: GCST007800, GCST90399721, GCST90244098). GWAS association scores per genomic location were mapped onto the lung gene regulatory network (eGRN), computed as described above. Snp2cell^72^ was then used to propagate^85^ association scores over the network and compute significant enrichments. Further, marker gene scores were computed for each cell type using differential expression, mapped onto the GRN, propagated and overlapped with disease association scores with snp2cell, to obtain cell-type specific disease associations.

---

**Extended Figure 1:**
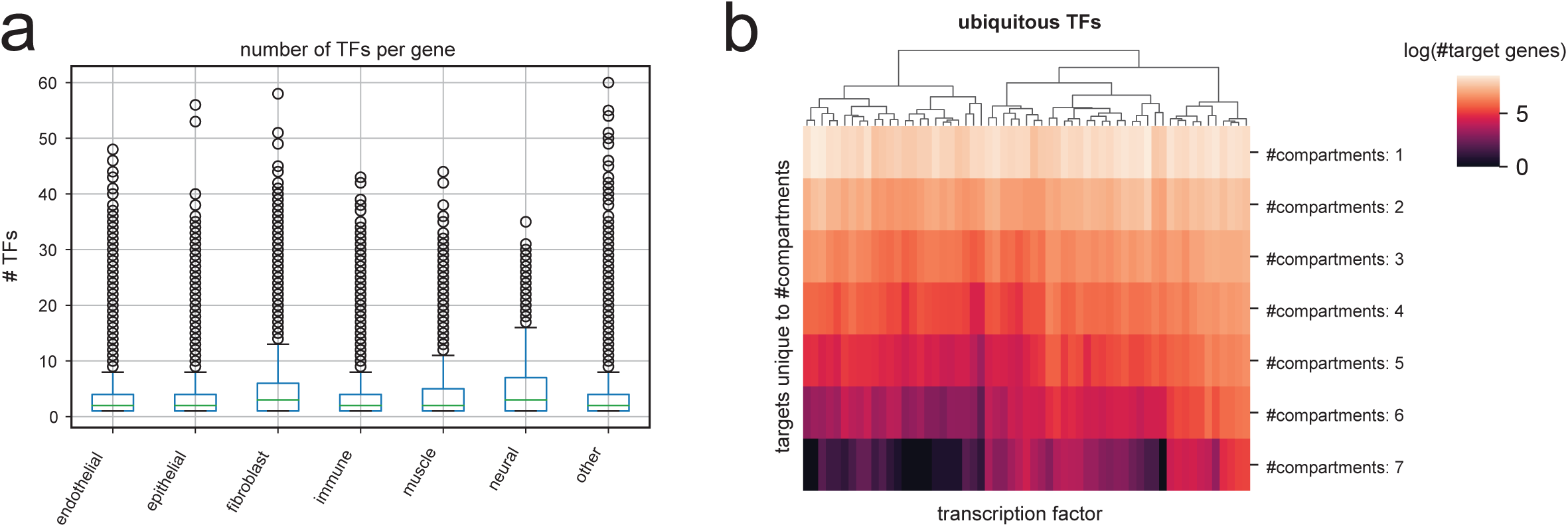
Regulon atlas overview. **(A)** Number of transcription factors regulating a single target gene, shown per cell compartment. Across compartments, most target genes are regulated by less than 5 TFs, while some are regulated by several dozen. **(B)** Heatmap of target gene numbers of ubiquitous TFs, showing by how many cell compartments they are shared (endothelial, epithelial, fibroblast, immune, muscle, neural, other). Although ubiquitous TFs are found across compartments, most of their target genes are found only in a small number of compartments, indicating compartment-specific functions.

**Extended Figure 3.**
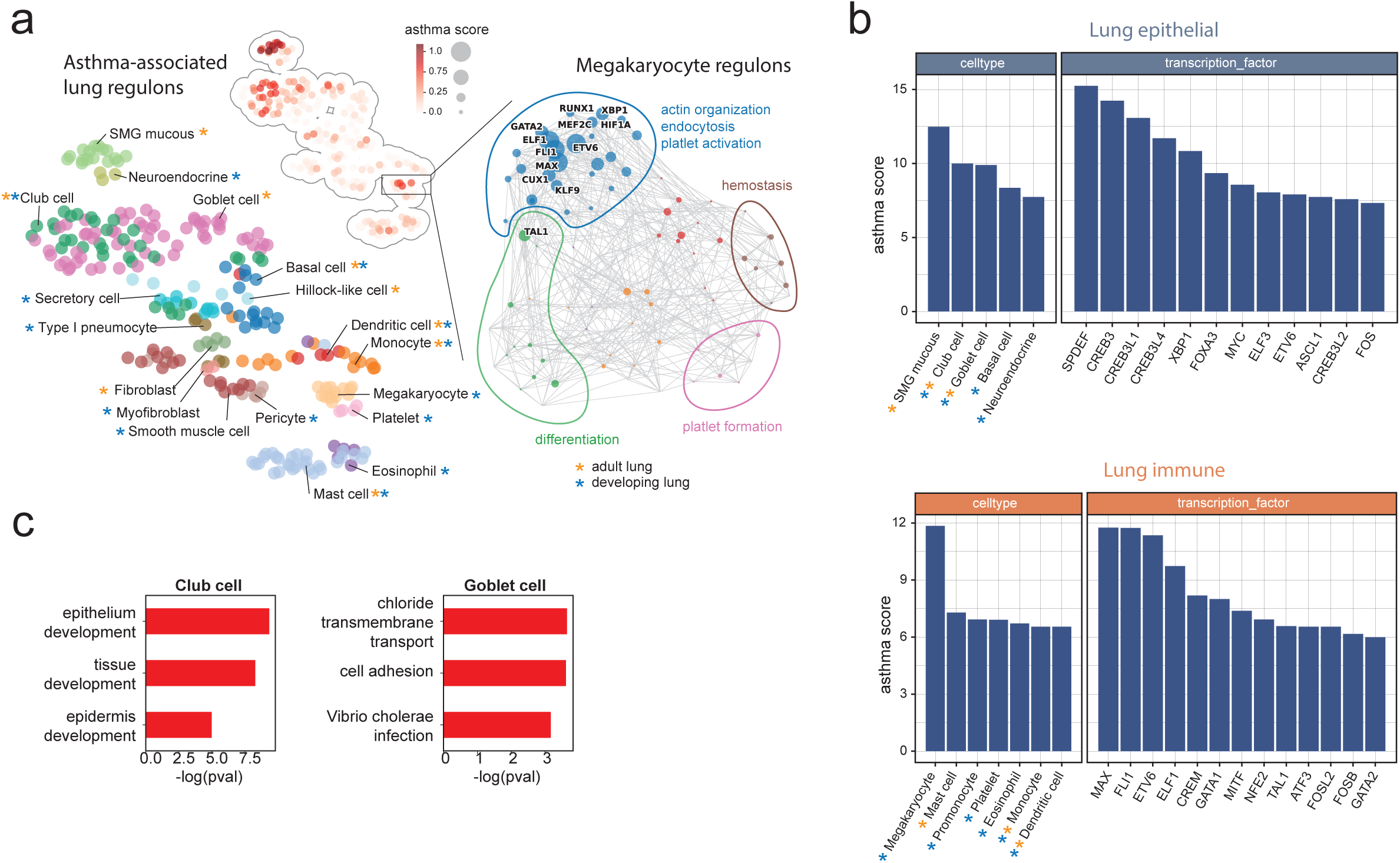
Asthma gene set association. **(A)** Regulon embedding based on target gene overlap, coloured by cell type, including regulons derived from adult and fetal lung data. Inset: asthma gene set scores. Right: Zoom-in into the regulon network for lung megakaryocytes. The node size is scaled by asthma score. Biological functions have been annotated manually. **(B)** Ranking of regulons for a consensus asthma gene set. Top: querying against lung epithelial cells. Bottom: querying against lung immune cells. **(C)** Pathway enrichment (g:profiler) of gene programs regulated by cell-type specific TF clusters with high asthma scores for club and goblet cells. Only genes unique to either club or goblet cells were used for the enrichment.

**Extended Figure 4.**
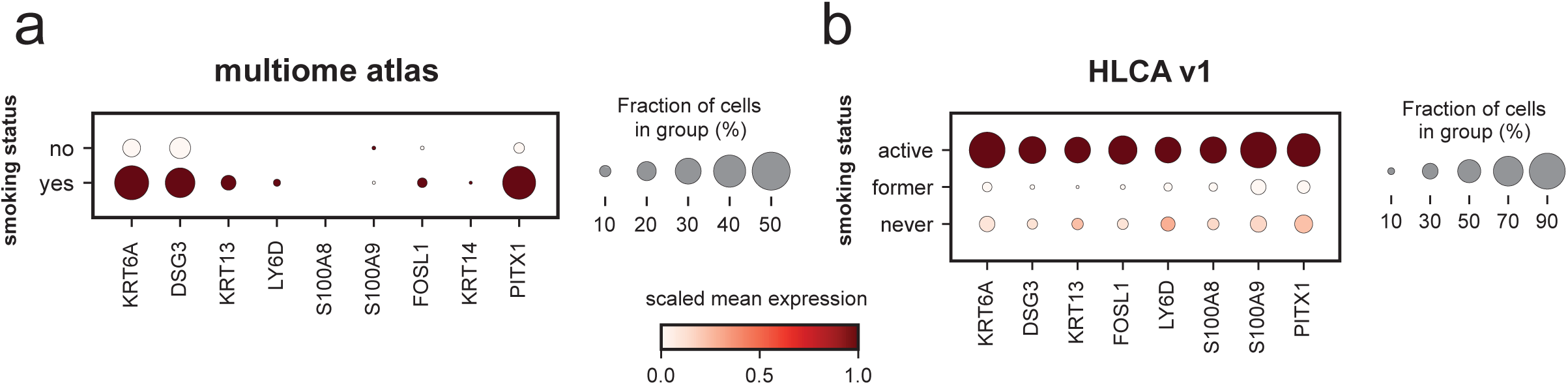
Hillock cells and smoking status. **(A)** Dotplot showing gene expression of hillock cell and keratinization markers in hillock and suprabasal cells of the lung multiome atlas, stratified by smoking status (5 donors). **(B)** Dotplot showing gene expression of hillock cell and keratinization markers in hillock and basal cells of the Human Lung Cell Atlas (HLCA) version 1, stratified by smoking status (57 donors with >10 cells).

**Extended Figure 5.**
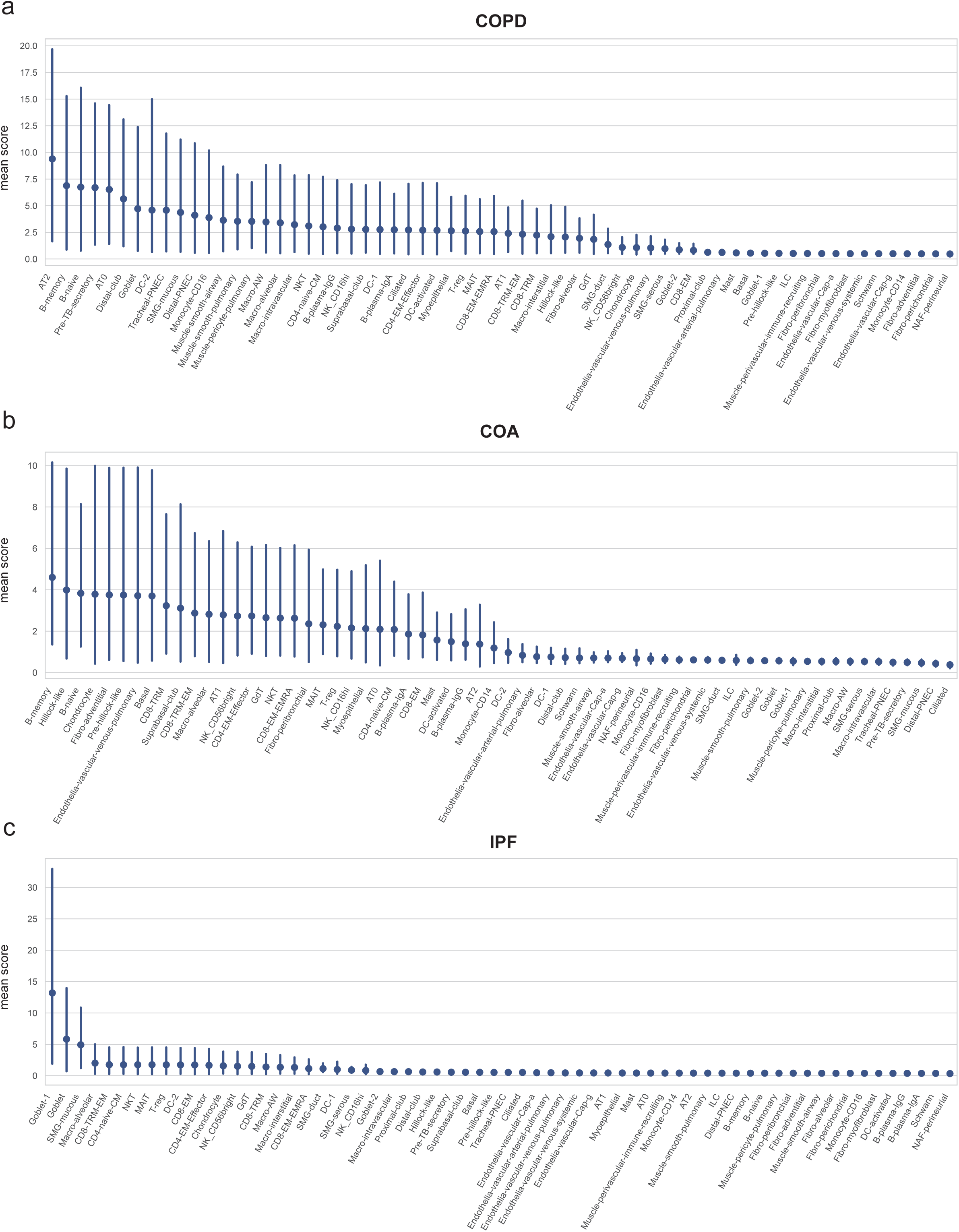
Genetic TF-associations in lung disease. Ranking of all annotated lung cell types based on average snp2cell scores for the top associated genes (z-score>10) for three diseases: **(A)** COPD, **(B)** childhood-onset asthma (COA), and **(C)** idiopathic pulmonary fibrosis (IPF).

